# Altered synaptic plasticity and central pattern generator dysfunction in a *Drosophila* model of PNKD3 paroxysmal dyskinesia

**DOI:** 10.1101/2020.02.20.957639

**Authors:** Simon Lowe, Patrick Kratschmer, James E.C. Jepson

## Abstract

**Background:** Paroxysmal non-kinesigenic dyskinesia type-3 (PNKD3) has been linked to gain-of-function (GOF) mutations in the hSlo1 BK potassium channel, in particular a dominant mutation (D434G) that enhances Ca^2+^-sensitivity. However, while BK channels play well-known roles in regulating neurotransmitter release, it is unclear whether the D434G mutation alters neurotransmission and synaptic plasticity in vivo. Furthermore, the subtypes of movement-regulating circuits impacted by this mutation are unknown.

**Objectives:** We aimed to use a larval *Drosophila* model of PNKD3 (*slo*^E366G/+^) to examine how BK channel GOF in dyskinesia alters synaptic properties and motor circuit function.

**Methods:** We used video-tracking to test for movement defects in *slo*^E366G/+^ larvae, and sharp-electrode recordings to assess the fidelity of Ca^2+^-dependent neurotransmitter release and short-term plasticity at the neuromuscular junction. We then combined sharp-electrode recording with ex vivo Ca^2+^-imaging to investigate the functionality of the central pattern generator (CPG) driving foraging behavior in *slo*^E366G/+^ larvae.

**Results:** We show that the PNKD3 mutation leads to Ca^2+^-dependent alterations in synaptic release and paired-pulse facilitation. Furthermore, we identify robust alterations in locomotor behaviors in *slo*^E366G/+^ larvae which were mirrored by dysfunction of the upstream, movement-generating CPG in the larval ventral nerve cord.

**Conclusion:** Our results demonstrate that a BK channel GOF mutation can alter neurotransmitter release and short-term synaptic plasticity, and result in CPG dysfunction, in *Drosophila* larvae. These data add to a growing body of work linking paroxysmal dyskinesias to aberrant neuronal excitability and synaptic plasticity in pre-motor circuits.

## Introduction

Paroxysmal dyskinesias (PxDs) are a heterogenous group of hyperkinetic movement disorders defined by the presence of intermittent dystonic, choreiform and/or ballistic involuntary movements^1^. Distinct forms of PxD can be clinically differentiated based on the triggers for attacks: paroxysmal kinesigenic dyskinesia (PKD; triggered by sudden movement), paroxysmal exercise-induced dyskinesia (PED), and paroxysmal non-kinesigenic dyskinesia (PNKD; often triggered by alcohol, caffeine, stress and fatigue)^2^. Here we focus on a PxD subtype whose neuropathological basis has not previously been investigated in an animal model: type-3 PNKD (PNKD3; OMIM #609446).

PNKD3 is characterised by dystonic or choreiform movements that can occur spontaneously or be triggered by alcohol, fatigue and stress^3 4^. Dyskinetic attacks emerge in early infancy or childhood, are frequently co-morbid with absence or generalized tonic-clonic seizures^3^, and in some cases occur alongside developmental delay^4^. PNKD3 is caused by dominant gain-of-function (GOF) mutations in *KCNMA1*, encoding the pore forming α-subunit of the hSlo1 BK (big K^+^ conductance) channel^3-7^. BK channels are voltage- and Ca^2+^-activated potassium channels, and modulate synaptic output from excitable cells by gating K^+^ efflux during the repolarization and afterhyperpolarization (AHP) phases of the action potential (AP)^8-10^. The most common PNKD3 mutation is a heterozygous missense mutation (1301A→G) in exon 10 of *KCNMA1*, resulting in the replacement of a negatively charged aspartic acid residue with a neutral glycine (D434G) in the cytoplasmic regulator of K^+^ conductance 1 (RCK1) domain^3^. RCK1 contains critical Ca^2+^-binding sites^11^, and in vitro evidence indicates that D434G acts as a gain-of-function mutation by increasing Ca^2+^ sensitivity, accelerating activation and decelerating deactivation^3 5 12^.

While the roles of BK channels in regulating AP dynamics and neurotransmitter release have been studied in many species^10^, it has remained unknown how the *KCNMA1*/hSlo1 D434G mutation impacts neurotransmission in vivo. This is a relevant issue, since recent studies have pointed towards alterations in synaptic transmission and short-term plasticity as a potential cellular phenotype disrupted across several PxD subtypes. For example, mutations in the *PRRT2* and *PNKD* loci cause PKD (OMIM #128200) and type-1 paroxysmal non-kinesigenic dyskinesia (PNKD1; OMIM #118800) respectively^13-17^. PRRT2 represses synaptic vesicle priming by inhibiting assembly of the pre-synaptic trans-SNARE complex^18 19^, and loss of PRRT2 results in increased synaptic facilitation during the later phases of repetitive stimuli at cerebellar Parallel Fibre-Purkinje Cell (PF-PC) synapses^19^. Likewise, overexpression of the PNKD protein in cultured hippocampal neurons suppresses neurotransmitter exocytosis^20^, and knock-out of *PNKD* enhances synaptic facilitation at hippocampal Schaffer collateral synapses^20^.

The precise circuits that drive involuntary movements in PNKD3 are also not defined. Studies from human patients and animal models have implicated central pre-motor neurons, including cerebellar and basal ganglia-thalamocortical circuits, in PxD pathogenesis^19 21 22^. However, peripheral circuits may also contribute to dyskinetic attacks. For example, abnormal spinal reciprocal inhibition has been identified in patients with PKD^23^, and secondary PxDs have been associated with spinal cord lesions, glioma and compression^24^. Defective sensory processing has also been observed in patients with focal and generalized dystonia and has been proposed to predispose towards dystonic motor phenotypes^25^. However, in the context of PNKD3, the lack of animal models has precluded tests of the relative contribution of these circuit types to motor dysfunction.

In our companion work we describe the generation of a *Drosophila* model of PNKD3 harbouring the equivalent mutation of hSlo1 D434G (E366G) in the highly conserved orthologue of *KCNMA1, slowpoke* (*slo*). We show that heterozygosity for the SLO E366G mutation causes PxD-like motor phenotypes and narrows APs in SLO-expressing neurons at the adult stage, validating the model and confirming predictions as to the effect of hSlo1 D434G on AP waveforms in native neurons^3^. Here we utilise the comparatively simple architecture of neural circuitry influencing movement in *Drosophila* larvae to test how the SLO E366G mutation impacts synaptic release, short-term plasticity and pre-motor circuit function.

Exploratory locomotor behaviour in *Drosophila* larvae follows a stereotyped pattern consisting of forward movements punctuated by sporadic turns^26^. Forward movement is generated by peristaltic waves of muscle contraction that sequentially spread from posterior to anterior segments of the larval body wall. These contractions are driven by rhythmic input to motoneurons from an intrinsically active central pattern generator (CPG) located in the thoracic and abdominal segments of the larval ventral nerve cord (VNC), which also initiates turning behavior^27^. Following muscle contractions, proprioceptive neurons in the larval body wall provide sensory feedback to pre-motor circuits, facilitating rapid propagation of the contractive wave from posterior to anterior segments and enhancing crawling speed^28^.

The function of pre-motor circuits and motoneurons driving larval locomotion can be interrogated via sharp electrode recordings and ex vivo calcium imaging^29-31^. Furthermore, the neuromuscular junction (NMJ) of *Drosophila* larvae represents a widely studied model glutamatergic synapse in which alterations in neurotransmitter release and/or synaptic plasticity can be readily assessed^32^. Taking advantage of these circuit characteristics, we demonstrate that BK channel GOF in the *Drosophila* PNKD3 model alters synaptic output and short-term plasticity in a Ca^2+^-dependent manner and strongly disrupts CPG function.

## Materials and methods

### *Drosophila* genetics and husbandry

Flies were reared on standard medium at 25°C under 12 hour light/dark cycles. Three control *slo*^*loxP*^ lines (132.1.1, 111.1.1 and 7.1.1) and experimental *slo*^*E366G*^ (25.1.1, 137.1.3 and 72.1.1) were generated as described in the companion publication. To control for background genetic differences, each recombinant allele was outcrossed for 5 generations into an isogenic control background, iso31, and balanced in trans over TM6b, *tb*. Experimental flies were generated by crossing these lines to iso31 to generate heterozygous lines referred to as *slo*^*loxP*/+^ and *slo*^E366G/+^. A recombinant of the *ok371*-Gal4 driver and the UAS-*GCaMP6m* Ca^2+^ indicator was a kind gift from Dr. Stefan Pulver (University of St. Andrews).

### Motor analysis

Age-matched ‘wandering’ third instar (L3) larvae were selected, gently cleaned with Milli-Q water, and transferred individually to a wide flat plane coated in 2%agar (Sigma-Aldrich), which was placed in an incubator (LSM) at 25°C and a relative humidity of 50-55%. For each larvae, after 10 s of acclimatization, 60 s of free movement was recorded using a Samsung S5 mobile phone at 15 fps and 640×480 resolution. Video-tracking was performed using the desktop version of the AnimApp video tracker^33^, available on https://github.com/sraorao/. A custom R script, which has been made available on GitHub (https://github.com/PatrickKratsch/), was used to calculate the following movement parameters: total distance travelled was calculated by summing the difference in XY position between consecutive frames; turns were identified by calculating changes in larval width:length ratio, with a threshold ratio of 0.4 manually identified as a reliable indicator of turning. There was no significant difference between the three control or experimental lines in either parameter: thus, the data were pooled. In subsequent electrophysiological and Ca^2+^ imaging experiments, one control (132.1.1) and one experimental (25.1.1) line were used.

### NMJ electrophysiology

#### Standard recordings

Sharp-electrode intramuscular voltage recordings were taken from muscle 6, abdominal segment 3 according to standard protocols^31 34^. Wandering L3 larvae were dissected in ice-cold, Ca^2+^-free modified HL3.1 saline^35^ consisting of 70 mM NaCl, 5 mM KCl, 10 mM NaHCO3, 115 mM sucrose, 5 mM trehalose, 5 mM HEPES, and 10 mM MgCl2. Motor nerves were severed just below the VNC, and the brain was removed. CaCl2 was added to the bath solution at the concentrations indicated. Recordings took place at 22-25°C. Sharp microelectrodes (thick-walled borosilicate glass capillaries, pulled on a Sutter Flaming/Brown P-97 micropipette puller) were filled with 3 M KCl and had resistances of 20–30 MΩ. Recordings were amplified using an Axon Instruments AxoClamp-2B, digitised at 25 kHz using a National Instruments DAQ, and analysed offline using StrathClyde Electrophysiology Software WinEDR v3.2.7. Input resistance was calculated by injecting a step current of 1 nA for 500 ms and deriving the resistance from the voltage change using Ohm’s law, with the pipette resistance subtracted by bridge balancing. Recordings were discarded if their initial RMP was more positive than −60mV, varied by >10%during recording, or if the input resistance was < 5 MΩ. EJPs were evoked by drawing the severed ends of motoneurons into a thin-walled glass suction electrode and stimulating with a single square-wave voltage pulse of 0.1 ms and 10 V (Digitimer DS3 Isolated Current Stimulator). To analyse EJP amplitude, motoneurons were stimulated 20 times at 5 s intervals, and the mean amplitude was taken. To analyse paired pulses, motoneurons were stimulated twice with a 0.1 s interval, and this was repeated 5 times at 10 s intervals. The amplitude of the second event was calculated as a percentage of the first, and the mean percentage was taken. Spontaneously occurring mEJPs were automatically identified using WinEDR during 150 s recording in the absence of stimulation, and verified manually. Mean mEJP amplitudes and inter-event intervals were taken from at least 100 events per recording.

#### Semi-intact recordings

The procedure was as above except the brain and motor nerves were left intact and care was taken not to damage them during dissection. To limit muscle contractions, 25 µM nifedipine (Sigma Aldrich) was added to the bath solution^36^. Nifedipine was pre-dissolved in DMSO, which was added to the solution with a final concentration of 0.2%. Each recording lasted 300 s in the absence of stimulation. Due to the high level of activity it was not possible to measure input resistance, so to ensure high fidelity recordings were discarded if the RMP did not remain more negative than −60 mV for the duration of recording. To analyse burst firing, a ‘burst’ was arbitrarily defined as starting when at least five EJPs occurred in succession at ≥5 Hz (i.e. 5 successive inter-EJP intervals of ≤ 0.2 s), and ending when at least 1 s passed with no EJPs occurring at >5 Hz (i.e. no inter-EJP intervals of ≤ 0.2 s).

### NMJ morphology

Immunostaining of larval NMJ was performed according to standard protocols^37^. Wandering L3 larva were dissected as above and fixed in 4%PFA for 10-20 mins at room temperature. Primary antibodies were goat anti-HRP conjugated Alexa Fluor 488 at 1:500 (Jackson ImmunoResearch) and mouse anti-DLG at 1:200 (clone 4F3, DSHB). Secondary antibody was goat anti-mouse Alexa Fluor 555 at 1:1000. NMJs innervating muscle 6/7, abdominal segment 3 were visualised using a Zeiss confocal LSM710 with an EC ‘Plan-Neofluar’ 20x/0.50 M27 air objective. Bouton number was counted manually and included both 1b and 1s type boutons. Bouton size, NMJ area and muscle area were quantified using ImageJ. All analyses were performed blind to genotype.

### GCaMP6m imaging

Age-matched L3 larvae of genotypes *ok371*-Gal4 >UAS-*GCaMP6m*, slo^E366G/+^ and *ok371*-Gal4 >UAS-*GCaMP6m*, slo^*loxP*/+^ were obtained by crossing *ok371*-Gal4, UAS-*GCaMP6m* recombinants to *slo*^*loxP*^/TM6b, *tb* or *slo*^E366G^/TM6b, *tb* flies, with non-*tb* 3^rd^ instar larvae subsequently selected for experiments. Larvae were dissected in a recording solution consisting of 135 mM NaCl, 5 mM KCl, 4 mM MgCl2, 2 mM CaCl2, 5 mM TES, and 36 mM sucrose^38^. The entire CNS was removed and immersed in the recording solution. Recordings were taken immediately after dissection on a Zeiss LSM 710 confocal microscope with an EC ‘Plan-Neofluar’ 20x/0.50 M27 air objective. Recordings lasted 5 min with a scan time of 390.98 ms per frame and 200 ms intervals between scans. Fluorescence activity was measured within regions of interest (ROIs) of consistent area (115.809 µm^2^) drawn around the dendritic regions of motoneurons in segments 7 and 4 on each side of the VNC. Background fluorescence was subtracted from each of these 4 datasets using an extra ROI drawn in the top left corner of the image lacking any neural tissue. Data from the four ROIs were plotted separately. Spikes in fluorescence were identified manually. ‘Forward waves’ were defined as concurrent spikes on both sides of segment 4 which were preceded by concurrent spikes on both sides of segment 7. ‘Forward wave ΔF’ was defined as the distance from peak to trough fluorescence of a segment 4 forward Ca^2+^ wave. ‘Frequency of forward wave’ was defined as the number of segment 4 forward wave peaks per second during each 5 min recording period. ‘Propagation time’ was defined as time between segment 7 and 4 peaks during a forward wave. ‘Turns’ were defined as asymmetric spikes occurring on only one side of segment 4.

## Results

### Disrupted locomotor patterns in *slo*^*E366G/+*^ larvae

In our accompanying article we describe a *Drosophila* model of PNKD3. We used ends-out homologous recombination to generate two distinct alleles of *slo*, the *Drosophila* ortholog of *KCNMA1*. One allele contains the equivalent mutation to hSlo1 D434G (E366G) and a *loxP* site in an upstream intron (Fig. 1A). The second allele contains the intronic *loxP* site without any exonic coding alterations, representing the control strain (Fig. 1A). Adult flies heterozygous for the E366G allele (*slo*^E366G/+^) exhibit reduced locomotion compared to controls as well as sporadic leg twitches. We were interested to explore whether PNKD3 model *Drosophila* also displayed motor deficits at an earlier stage of the *Drosophila* lifecycle: the 3^rd^ instar (L3) larval stage. We assessed larval motor behaviour in *slo*^E366G/+^ and *slo*^*loxP*/+^ L3 larvae during 1 min of video-tracked free movement on a flat agar plane (Fig. 1B, C). Both the total distance moved and number of turns initiated by *slo*^E366G/+^ larvae were substantially reduced relative to *slo*^*loxP*/+^ controls (Fig. 1D, E). Thus, the *slo*^E366G^ mutation induces a robust perturbation of motor control at the larval stage of the *Drosophila* life cycle.

**FIG 1.**
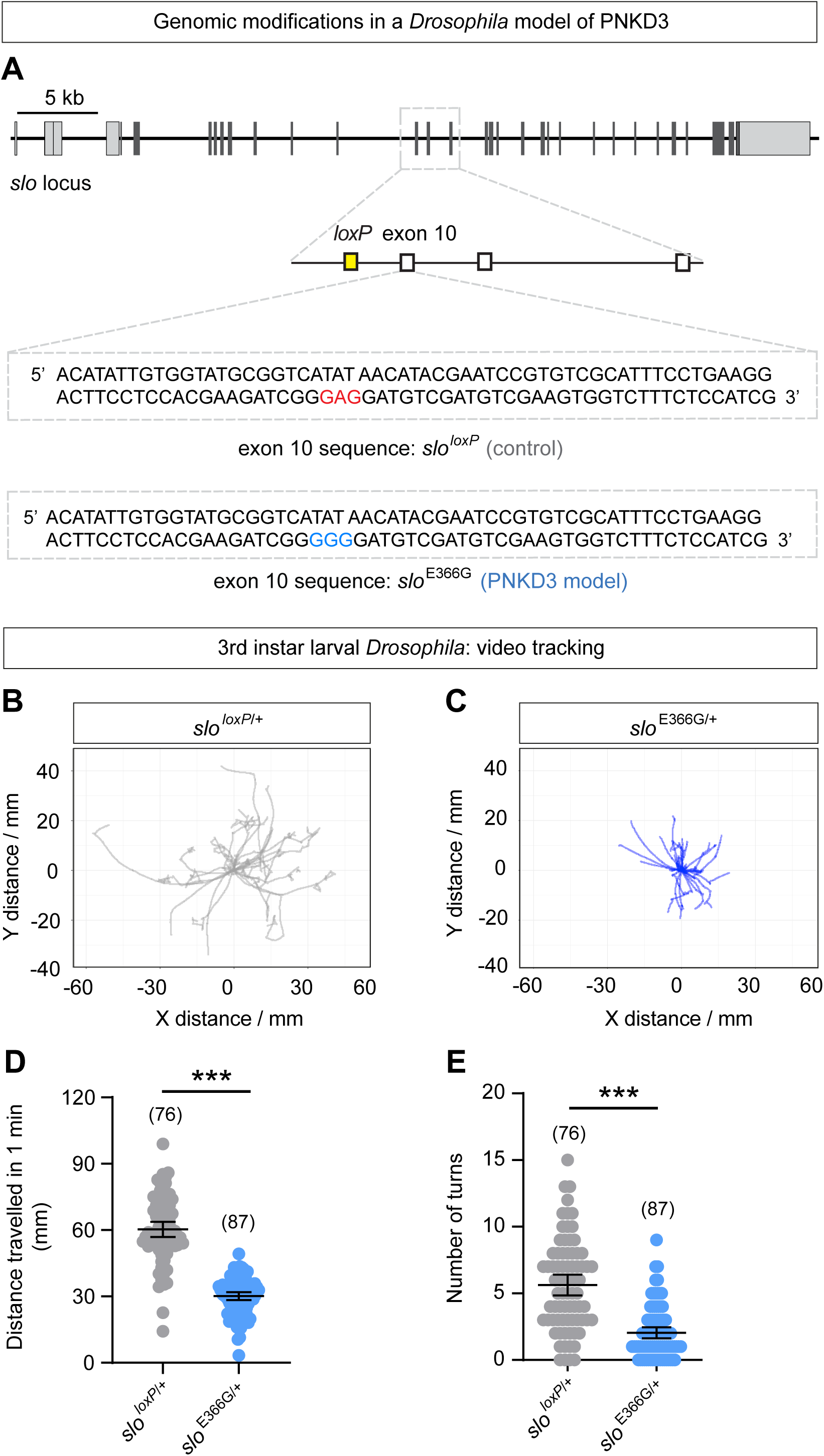
(A) Schematic illustrating genomic modifications in the PNKD3 *Drosophila* model *slo*^E366G/+^ and associated control (*slo*^*loxP*/+^) strains. Light grey boxes: 5’ and 3’ untranslated regions. Dark grey boxes: coding exons. The sequence of exon 10, containing the E366 residue, is shown below. In the *Drosophila* PNKD3 model, the GAG codon encoding E366 has been mutated to GGG, resulting in the E366G mutation. (B-C) Example overlaid traces of paths travelled by individual *slo*^E366G/+^ and *slo*^*loxP*/+^ L3 larvae during 1 minute free movement on agar plane. Larvae were derived from single *slo*^E366G/+^ and *slo*^*loxP*/+^ recombinant lines. (D-E) Mean distance travelled (D) and number of turns (E) over 1 min of *slo*^E366G/+^ and *slo*^*loxP*/+^ L3 larvae. Data is pooled from 3 independently generated lines for each genotype. Dots represent individual larvae; n-values are noted. Error bars: mean and 95% Confidence Interval (CI). ***p<0.0005, unpaired t-test with Welch’s correction (D) or Mann-Whitney U-test (E).

### Altered Ca^2+^-dependent neurotransmitter release at the neuromuscular junction of *slo*^*E366G/+*^ larvae

While the reduction in turning behavior in *slo*^E366G/+^ larvae is suggestive of CPG dysfunction^27^, defective forward peristalsis could also arise from reduced neurotransmitter release at the neuromuscular junction (NMJ). Indeed, loss of SLO channels has previously been shown to enhance neurotransmitter release from, and alter the development of, motoneuron synaptic boutons^39^. We therefore examined the morphology and physiological properties of the NMJ in *slo*^E366G/+^ and *slo*^*loxP*/+^ larvae.

Neither the passive membrane properties of the muscle (Supplemental Fig. 1) nor the morphology of motoneuron synapses (Supplemental Fig. 2) was significantly different between *slo*^E366G/+^ and *slo*^*loxP*/+^ L3 larvae. We therefore next tested for alterations in evoked neurotransmission. We examined evoked excitatory junction potentials (EJPs) using sharp electrode recordings (Fig. 2A). Since the hSlo1 D434G mutation enhances BK channel Ca^2+^-sensitivity^7^, we measured EJP amplitudes over a range of extracellular Ca^2+^ concentrations ([Ca^2+^]_e_; 0.15 – 3 mM) (Fig. 2B-F). Consistent with previous work^40^, we observed a strong positive correlation between [Ca^2+^]_e_ and EJP amplitude in *slo*^E366G/+^ and *slo*^*loxP*/+^ NMJs (Fig. 2B-F). Furthermore, in support of an enhancement of BK channel Ca^2+^-sensitivity, *slo*^E366G/+^ EJPs were significantly smaller compared to controls at lower [Ca^2+^]_e_, but were unchanged at higher [Ca^2+^]_e_ (Fig 2B-F).

**FIG 2.**
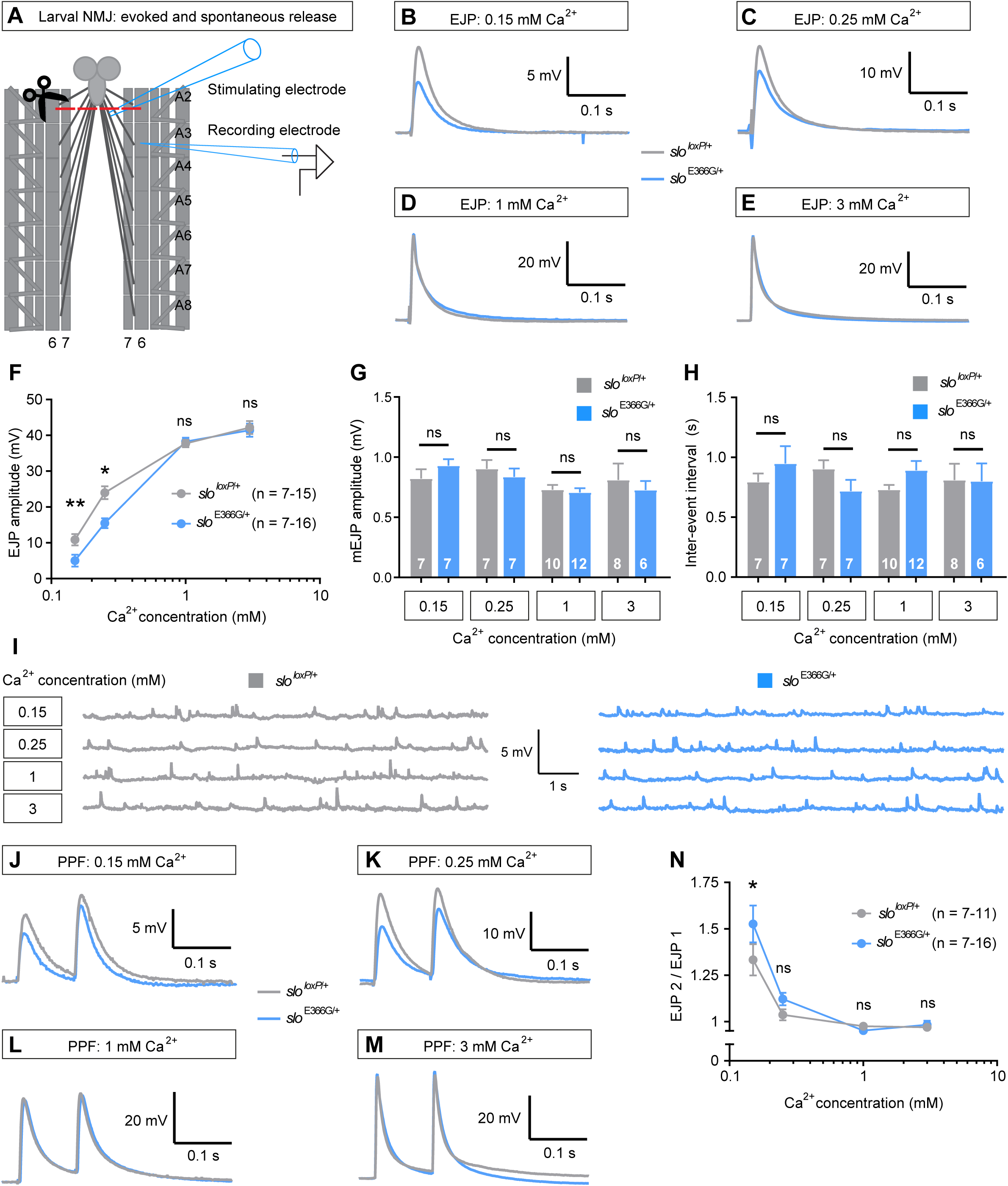
(A) Illustration of the electrophysiological protocol used. A sharp intramuscular recording electrode records from segment A3 of the longitudinal body wall muscle 6. Motoneurons innervating the body wall muscles are severed just below the VNC and EJPs are evoked by stimulating the severed end of the motoneurons innervating muscle 6 A3. Abdominal segments A2-A8 are shown. (B-E) Representative EJP waveforms from *slo*^E366G/+^ and *slo*^*loxP*/+^ larvae across a range of [Ca^2+^]_e_. (F) EJP amplitudes at each [Ca^2+^]_e_ (X axis shown as log_10_). (G-H) mEJP amplitude (G) and inter-event interval (H) at each [Ca^2+^]_e_. (I) Representative mEJPs from *slo*^E366G/+^ and *slo*^*loxP*/+^ larvae across a range of [Ca^2+^]_e_. (J-M) Representative paired-pulse waveforms at each [Ca^2+^]_e_. (N) Paired-pulse ratio shown as EJP2 / EJP1 at each [Ca^2+^]_e_ (X axis shown as log_10_). n-values are noted. Error bars: mean ± SEM. *p<0.05, **p<0.005 ***p<0.0005, ns – p>0.05, two-way ANOVA with Sidak’s multiple comparisons test (F, M) or Mann-Whitney U-test (G, H).

The Ca^2+^-dependent alteration in synaptic transmission at the NMJ of *slo*^E366G/+^ larvae was not accompanied by alterations in the amplitude or frequency of spontaneously occurring miniature EJPs (mEJPs) (Fig. 2G-I), suggesting a pre-synaptic locus for this effect. To provide support for this premise, we tested for alterations in short-term plasticity in *slo*^E366G/+^ larvae over the same range of [Ca^2+^]_e_. The relationship between [Ca^2+^]_e_ and short-term plasticity is well established at this synapse. Low [Ca^2+^]_e_ results in lower vesicle release per AP, smaller resulting EJPs, and paired-pulse facilitation (PPF). Conversely, increasing [Ca^2+^]_e_ enhances vesicle fusion, increases EJP amplitudes, and reduces PPF^41 42^. We found that *slo*^E366G/+^ larvae displayed a significant increase in PPF at 0.15 mM Ca^2+^, a non-significant trend towards an increase at 0.25 mM Ca^2+^, and no alteration at 1 or 3 mM Ca^2+^ (Fig. 2J-N), consistent with a reduction in neurotransmitter release at low [Ca^2+^]_e_ in *slo*^E366G/+^ larvae.

Collectively, these results demonstrate that increased presynaptic BK channel function due to the SLO 366G mutation (and by extension, hSlo1 D434G^3^) can reduce neurotransmitter release under particular physiological conditions (see Discussion). The higher [Ca^2+^]_e_ tested, in which EJP amplitude was unaltered in the *slo*^E366G/+^ background (Fig. 2F), are closer to the physiological composition of the larval haemolymph^43^, suggesting that intrinsic motoneuron function is largely unimpeded in *slo*^E366G/+^ larvae. These results point to a non-motoneuron origin of altered foraging locomotor behaviour in *slo*^E366G/+^ larvae. Since prior work has shown that activity of the CPG in the ventral nerve cord (in the absence of higher brain function or sensory feedback) is sufficient to drive larval peristalsis and turning behaviors^27^, we next sought to assess CPG activity in *slo*^E366G/+^ larvae.

### Perturbed rhythmicity of spontaneous motoneuron firing in *slo*^*E366G/+*^ larvae

To investigate CPG function in *slo*^E366G/+^ larvae we recorded spontaneous patterns of CPG-driven motoneuron activity by measuring intramuscular voltage changes in preparations in which the central and peripheral nervous systems are left intact. Spontaneous, rhythmic bursts of motoneuron activity can be readily observed in such preparations, which correlate with contraction of the innervated muscle segment in a behaving animal^44^ (Fig. 3A). Importantly, proprioceptive feedback to pre-motor circuits can be largely suppressed via inhibition of muscle contraction by the VGCC blocker nifedipine^36^, allowing the functionality of the CPG to be assessed in relative isolation.

**FIG 3.**
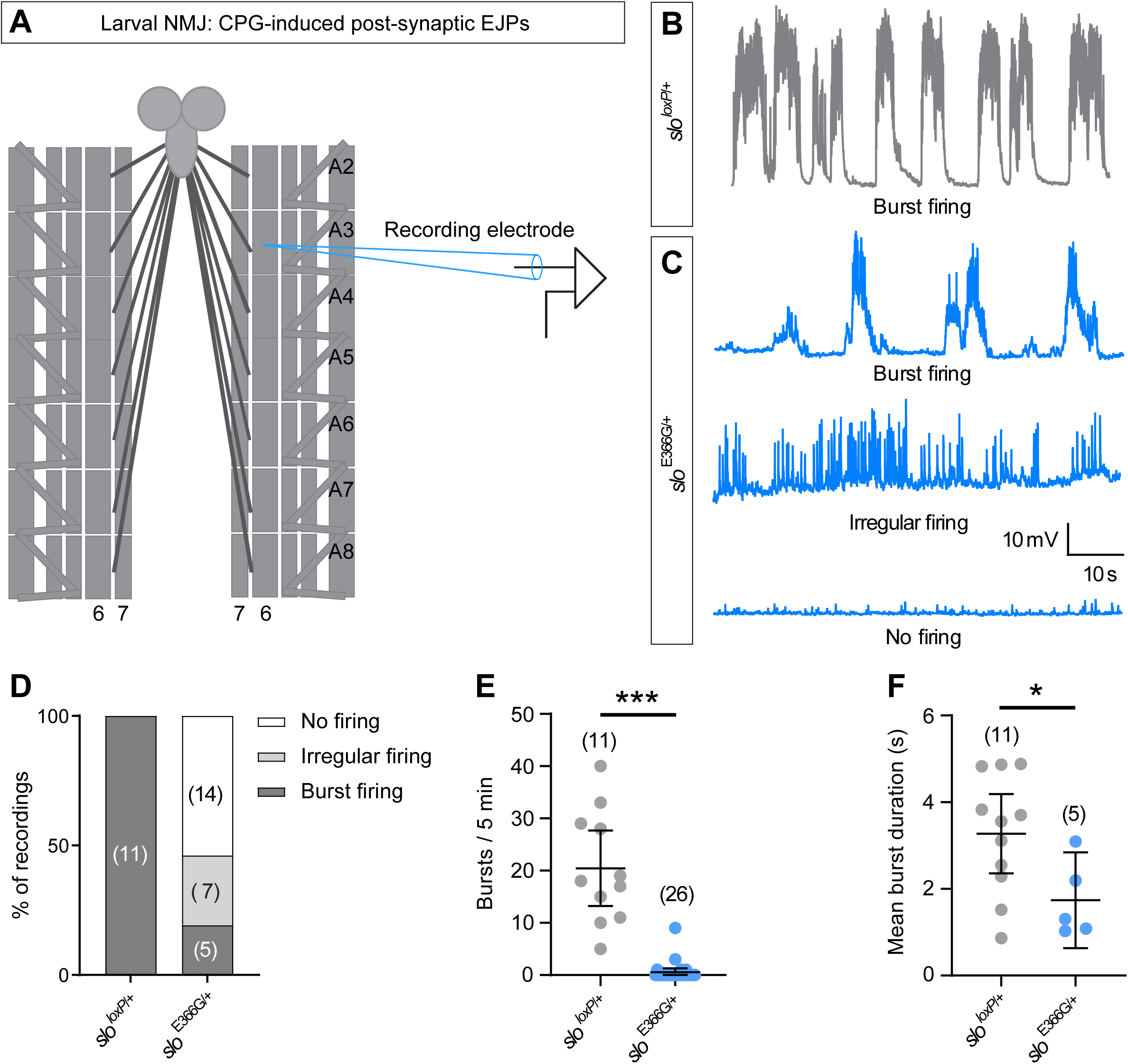
(A) Illustration of the electrophysiological protocol used. In contrast to the analysis of evoked EJPs (Fig. 2A), motoneuron axons innervating the body wall are left intact. Post-synaptic EJPs are thus elicited via activation of motoneurons by the upstream larval CPG. (B) Representative trace of spontaneous burst firing in a *slo*^*loxP*/+^ larva. (C) Representative traces of three distinct patterns of firing observed in *slo*^E366G/+^ larvae: burst firing, irregular firing, and no firing. (D) Percentage of *slo*^*loxP*/+^ and *slo*^E366G/+^ larvae showing each firing pattern. (E) Number of bursts in 5 minutes of recording. (F) Duration of bursts, including only recordings in which at least one burst occurred. Dots represent individual larvae; n-values are noted. Error bars: mean and 95% Confidence Interval (CI). *p<0.05, ***p<0.0005, Mann-Whitney U-test (E) or unpaired t-test with Welch’s correction (F).

Consistent with previous work^31 44^, we observed periodic bursts of high-frequency firing at regular intervals in all *slo*^*loxP*/+^ control larvae (Fig. 3B, D). Similar bursts were observed in only a minority (5/26) of *slo*^E366G/+^ larvae, and where they did occur were less frequent and of shorter duration (Fig. 3C-F). Interestingly, these data are mirrored by prior work showing that reducing SLO expression enhances the frequency of CPG-induced motoneuron bursts^29^. Thus, loss and gain of BK channel function results in bidirectional alterations in the endogenous motoneuron firing in *Drosophila* larvae. The majority of *slo*^E366G/+^ larvae (14/26), however, showed no firing activity during 5 minutes of recording, while the remainder (7/26) showed continuous, irregular activity that was not co-ordinated into bursts and quiescent periods (Fig 3C-F). Therefore, although largely functional (Fig. 2D-F), motoneurons in *slo*^E366G/+^ larvae display abnormal patterns of spontaneous activity, pointing towards dysfunction of the movement-driving CPG in *slo*^E366G/+^ larvae.

### The PNKD3 mutation disrupts CPG activity in the larval ventral nerve cord

Peristaltic larval locomotion requires co-ordinated alterations in activity across a population of motoneurons located in the VNC. While the precise circuit structure of the CPG driving rhythmic oscillations in motoneuron activity has yet to be fully defined, prior work indicates that the core CPG is also located in the larval VNC^27^. Importantly, CPG function persist in ex vivo preparations in the absence of either sensory feedback or input from the central brain^27 45^.

To investigate the extent of disruption to CPG activity in *slo*^E366G/+^ larvae, we visualised CPG-driven excitatory input to motoneuron dendrites in ex vivo larval brains. To do so, we expressed a genetically encoded fluorescent Ca^2+^ sensor, GCaMP6m^46^, under the *ok371*-Gal4 driver, which labels glutamatergic motoneurons in the VNC^47^ (Fig. 4A). Under live fluorescent imaging, motoneuron cell bodies and associated dendritic fields expressing GCaMP6m were clearly identifiable in the abdominal segments the VNC (Fig. 4A-C). Fictive forward locomotor patterns were visible as rhythmic waves of increased GCaMP6m fluorescence that moved from posterior to anterior motoneuron dendrites and cell bodies and were largely coincident between motoneuron fields in the left and right hemispheres (Fig. 4B, C and Supplemental Video 1). We detected numerous alterations in the pattern of CPG-driven fictive locomotor activity in *slo*^E366G/+^ motoneurons. Ca^2+^ waves moving from posterior to anterior were reduced in both amplitude and frequency in *slo*^E366G/+^ VNCs (Fig 4B-E), while the propagation of these waves from segments A7-4 was more rapid in the *slo*^E366G/+^ background (Fig 4F). It was also possible to identify fictive turns. Dendritic Ca^2+^ spikes were usually concurrent between motoneurons located in the left and right sides of the VNC. However, as described previously^30^, occasionally a spike occurred alongside a coincident trough in GCaMP6m fluorescence in the contralateral motoneuron dendrite (Fig. 4B). In behaving larvae this would result in muscle contraction on one side of the body, facilitating a turn. These fictive turns also occurred less frequently in *slo*^E366G/+^ VNCs (Fig. 4G).

**FIG 4.**
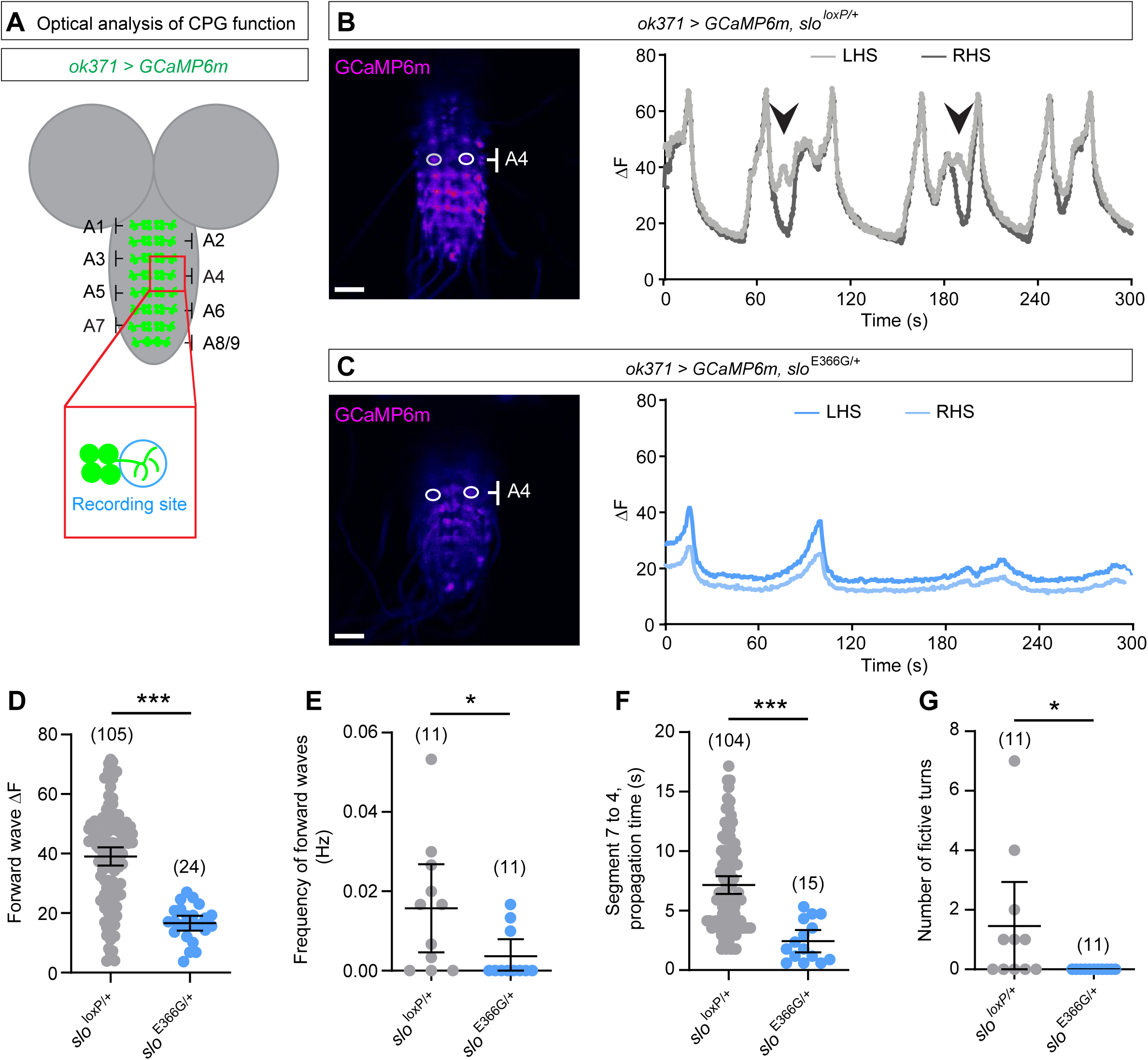
(A) Illustration of GCaMP6m-labelled motoneuron cell bodies and dendritic regions in the VNC and location of recording area around motoneurons in the abdominal A4 segment of the VNC. (B-C) Representative images showing GCaMP6m-labelled motoneuron cell bodies and dendrites in the VNC, and location of ROIs drawn around motoneuron dendrites on the left-and right-hand side of the A4 segment (LHS and RHS, respectively). Representative traces of fluorescence change in LHS and RHS over 300 s in *slo*^loxP/+^ and *slo*^E366G/+^ are shown. Arrowheads indicate fictive turns, where the LHS and RHS motoneuron dendrites exhibit opposing patterns of excitation. (D-G) Parameters of fictive locomotor patterns. Dots represent individual larvae; n-values are noted. Error bars: mean and 95% Confidence Interval (CI). *p<0.05, ***p<0.0005, Mann-Whitney U-test.

These data demonstrate abnormal patterning of motoneuron firing, with altered frequency, amplitude and co-ordination of intrinsically generated fictive motor behaviours, closely mirroring the changes in locomotor behaviour observed in foraging *slo*^E366G/+^ larvae (Fig. 1B-E). Given our electrophysiological evidence that motoneurons appear functionally intact under physiological conditions, we conclude that the PNKD3-associated SLO E366G mutation dominantly disrupts pre-motor circuit function, in particular the larval CPG, whose input into motoneurons is required for co-ordinated locomotor behaviour.

## Discussion

PNKD3 has been linked to GOF mutations in the hSlo1 BK channel^3-7 12^. BK channels are widely expressed throughout the mammalian nervous system^48^ and modulate numerous aspects of neuronal physiology, including axonal AP dynamics and presynaptic neurotransmitter release^10^. Yet due to the lack of in vivo models of PNKD3, how pathologically enhanced BK channel activity in PNKD3 impacts synaptic output and sensorimotor circuit activity has remained unclear. Here, focusing on the larval stage of the *Drosophila* life cycle, we demonstrate that the equivalent mutation to hSlo1 D434G in *Drosophila* (SLO 366G) results in Ca^2+^-dependent alterations in synaptic release and short-term plasticity, dysfunction of the larval locomotor CPG, and corresponding alterations in foraging behavior.

The hSlo1 D434G mutation has been theorized to increase excitability of pre-motor circuits by narrowing AP duration, thus accelerating the recovery of sodium channels from inactivation and enhancing AP frequency^3^. While we cannot rule out such an effect, in our *Drosophila* model we observed a suppression of neurotransmitter release at low [Ca^2+^]_e_ at the larval NMJ. Furthermore, the frequency of CPG-driven excitatory input to motoneuron dendrites is reduced in *slo*^E366G/+^ larvae. Together, these data suggest that the SLO E366G mutation may reduce excitability in neural circuits within the larval CPG, resulting in locomotor dysfunction. This postulate is in agreement with previously defined roles of BK channels in both vertebrate and invertebrate synapses. For example, at rat hippocampal CA3-CA3 synapses and the NMJs of nematodes and frogs, BK channels reduce release probability and synaptic output^49-52^, likely by narrowing AP duration and driving inactivation of presynaptic VGCCs required for Ca^2+^-dependent synaptic vesicle fusion^10^. Our data suggest that at the *Drosophila* NMJ, neurotransmitter release would be affected in *slo*^E366G/+^ larvae only under non-physiologically low [Ca^2+^]_e_. This may be due to the large safety factor and presence of numerous homeostatic mechanisms that ensure consistency of release at this synapse^53-55^. We therefore hypothesize that the SLO E366G mutation, and by extension hSlo1 D434G, primarily impacts central synapses that facilitate more graded, analogue changes in synaptic output. In particular, we posit that due to the enhanced Ca^2+^-sensitivity of E366G/D434G BK channels^7^, their impact may be highest at synapses where the coupling between BK channels and VGCCs is comparatively weak.

Such differences in the degree of coupling can occur via several mechanisms, including synapse-specific association of BK channels with distinct VGCC subtypes^56 57^, and variations in the degree of Ca^2+^ buffering within the VGCC Ca^2+^ nanodomain^58^. Combined with prior results demonstrating that the pathogenic effects of D434G on hSlo1 channel function are modified by auxiliary BK channel β-subinits^12^, each of which has a distinct expression pattern, these data begin to facilitate an understanding of which neural circuits may be most affected in PNKD3. While we cannot rule out any changes in proprioceptive signalling in our *Drosophila* model, our data strongly suggests that dysfunction of the pre-motor CPG is sufficient to explain the reduction in peristaltic and turning behavior during foraging observed in *slo*^E366G/+^ larvae. Thus, while altered activity with cerebellar and basal ganglia-thalamocortical circuits appears a common hallmark of PxDs^19 21 22^, in future mammalian models of PNKD3 it will also be of interest to test for dysfunction of spinal CPG networks that transform descending command signals from brain regions such as the mesencephalic locomotor region into coordinated movements^59 60^.

Further investigations are required to elucidate why the larval CPG in *Drosophila* is so strongly perturbed by BK channel GOF. Experimental and theoretical work have suggested that synaptic depression is important for maintaining phase invariance between neural components of CPGs^61-63^. Given the enhanced PPF at the NMJ of *slo*^E366G/+^ larvae under low [Ca^2+^]_e_, it is tempting to speculate that BK channel GOF may reduce the degree of synaptic depression or lead to a switch towards synaptic facilitation in key CPG circuits, resulting in a breakdown of phase-locked activity between CPG components. Continuing insights into the structure of larval pre-motor circuits^64^ may facilitate future investigations of this hypothesis.

Finally, in the context of paroxysmal non-kinesigenic dyskinesias it is interesting to note that the PNKD and hSlo1 proteins (linked to PNKD1 and PNKD3 respectively) may be components of a common protein interaction network within pre-synaptic termini. PNKD interacts and stabilizes the scaffold protein RIM1α, which promotes vesicle priming, docking and VGCC localization at the active zone^65 66^. RIM1αis itself bound by a family of RIM-binding proteins (RBPs) which ensure high-fidelity neurotransmission in cultured hippocampal neurons and at the Calyx of Held, likely by tethering VGCCs in close proximity to the active zone^67^. Intriguingly, RBPs also associate with BK channels and promote their coupling to presynaptic VGCCs in mammalian neurons^68^. In concert with these findings, the data presented here add support to the hypothesis that perturbations in a presynaptic protein interaction network influencing neurotransmitter release and/or Ca^2+^-dependent plasticity in pre-motor circuits, and potentially downstream movement-controlling CPGs, drives PNKD pathogenesis^20^.

## Acknowledgements

We thank Dr. Stefan Pulver (University of St. Andrews) for *ok371*-Gal4, UAS-*GCaMP6m* flies; Prof. Kirill Volynski (University College London), Dr. Edgar Buhl (Bristol University), and Prof. James Hodge (Bristol University) for comments on the manuscript; Dr. Srinivasa Rao Rao (University of Oxford) for help with larval video-tracking analyses; and Tom Lowe for help with video editing.

## Author Contributions

Conceptualization: S.L., P.K., J.E.C.J. Methodology: S.L., P.K., J.E.C.J. Software: P.K. Validation: S.L., P.K., J.E.C.J. Formal Analysis: S.L., P.K., J.E.C.J. Investigation: S.L., P.K., J.E.C.J. Writing – Original Draft: J.E.C.J. Writing – Review and Editing: S.L., P.K., J.E.C.J. Visualisation: S.L., P.K., J.E.C.J. Supervision: J.E.C.J. Project Administration: J.E.C.J. Funding Acquisition: P.K., J.E.C.J.

## Supplemental Figures, Figure Legends and Video Legends

**Supplemental Fig. 1.**
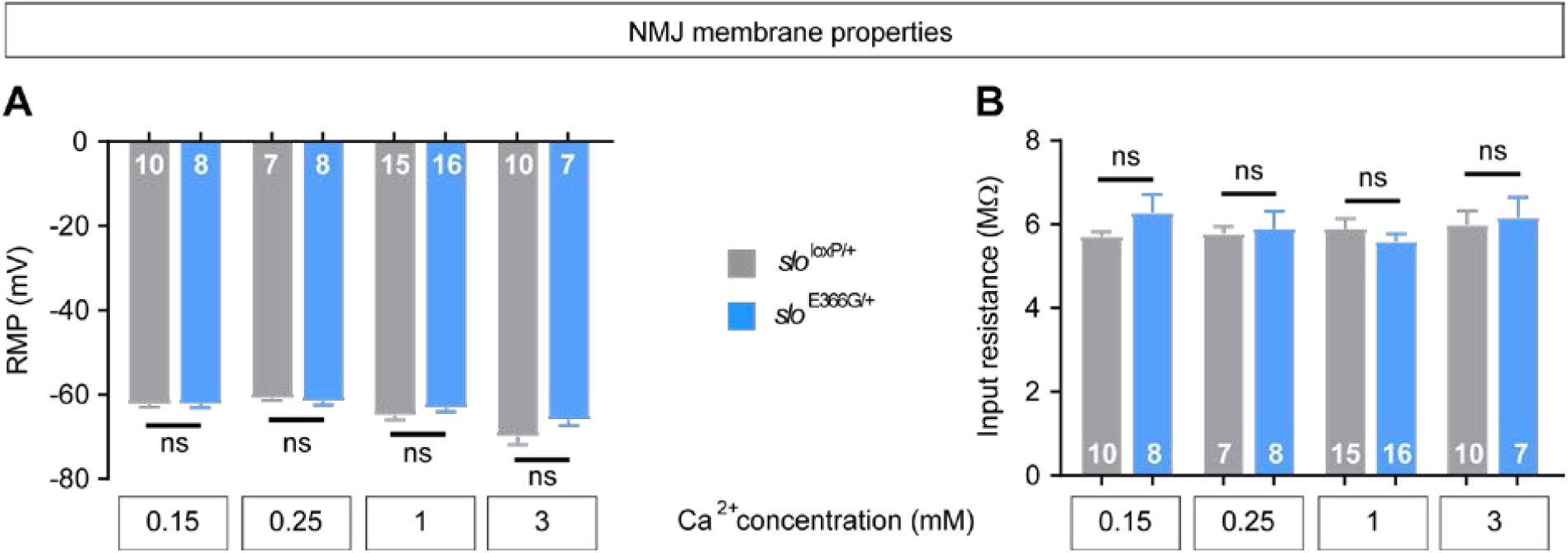
(A) Resting membrane potential (RMP) of muscle 6 in slo^*loxP*^/+ and slo^E366G/+^ larvae at varying [Ca^2+^]_e_. (B) Muscle input resistance in the same genotypes and [Ca^2+^]_e_. n-values are noted. Error bars: mean and SEM. ns – p>0.05, Mann-Whitney U-test (A. 0.25 mM and 1 mM Ca^2+^) or unpaired t-test with Welch’s correction (all other comparisons).

**Supplemental Fig. 2.**
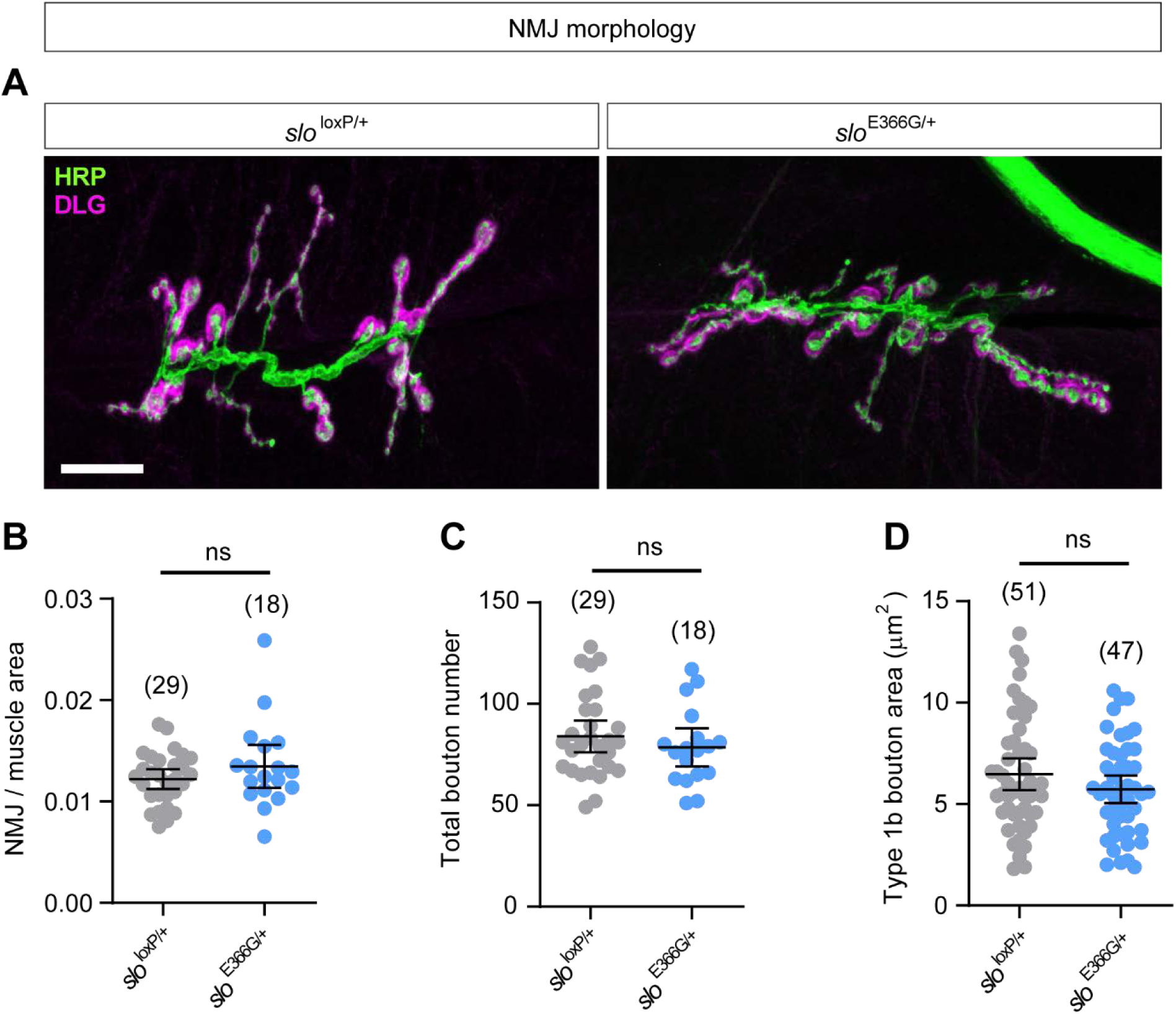
(A) Representative confocal images of HRP-labelled motoneurons innervating muscle 6/7, segment 3 of the 3^rd^ instar larval body wall. The post-synaptic sub-synaptic reticulum is labelled with anti-Discs Large (DLG). Scale bar: 20 µm. (B) NMJ area of slo^*loxP*^/+ and slo^E366G/+^ larvae normalized to the corresponding area of muscle 6/7. (C) Total bouton number (type 1s and type 1b) of motoneurons innervating muscle 6/7 in slo^*loxP*^/+ and slo^E366G/+^ larvae. (D) Area of type 1b boutons in slo^*loxP*^/+ and slo^E366G/+^ larvae. n-values are noted. Dots represent measurements of NMJ size/bouton number from individual larvae (B, C) or from individual synaptic boutons (D). Error bars: mean and 95%Confidence Interval (CI). ns – p>0.05, Mann-Whitney U-test (A) or unpaired t-test with Welch’s correction (B, C).

**Supplemental Video 1**. Video shows representative examples of waves of motoneuron excitation in the ex vivo larval ventral nerve cord (VNC) visualised via GCaMP6m driven by the glutamatergic neuron driver *ok371*-Ga4. In the *slo*^*loxP*/+^ control VNC (left), repetitive waves of excitation traveling from posterior to anterior segments can clearly be observed. Unilateral motoneuron excitation in the upper segments can also be infrequently observed. In the *slo*^E366G/+^ VNC (right), both the magnitude and frequency of increases in motoneuron GCaMP6m fluorescence is reduced, indicative of perturbed input from the upstream central pattern generator.

